# Targeting the Administration of Ecdysterone in Doping Control Samples

**DOI:** 10.1101/685230

**Authors:** Maria Kristina Parr, Gabriella Ambrosio, Bernhard Wuest, Monica Mazzarino, Xavier de la Torre, Francesca Sibilia, Jan Felix Joseph, Patrick Diel, Francesco Botrè

**Affiliations:** Institute of Pharmacy, Freie Universität Berlin, Germany; Agilent Technologies, Santa Clara CA, USA; Laboratorio Antidoping FMSI, Rome, Italy; Core Facility BioSupraMol, Department of Biology, Chemistry, Pharmacy, Freie Universitaet Berlin, Germany; Department for Molecular and Cellular Sports Medicine, Institute for Cardiovascular Research and Sports Medicine, German Sport University Cologne, Cologne, Germany; Department of Experimental Medicine, ‘Sapienza’ University of Rome, Italy

**Keywords:** Doping control, monitoring program, ecdysterone, urine analysis, LC-MS/MS, GC-MS, accurate mass

## Abstract

**Purpose:** The phytosteroid ecdysterone was recently reported to enhance performance in sports and may thus be considered as a substance of relevance in anti-doping control. To trace back an administration of ecdysterone from urine samples analytical properties have been investigated to assess its integration into initial testing procedures (ITP) in doping control laboratories.

**Methods:** Analytical properties of ecdysterone were evaluated using GC-QTOF-MS and LC-QTOF-MS. Its metabolism and elimination in human were studied using urines collected after administration.

**Results:** The detectability of ecdysterone by GC-MS (after derivatization) and/or LC-MS(/MS) has been demonstrated and sample preparation methods were evaluated. Dilute-and-inject for LC-MS(/MS) or SPE using Oasis HLB for GC-MS or LC-MS were found most suitable, while liquid-liquid extraction was hampered by the high polarity of ecdysteroids.

Most abundantly, ecdysterone was detected in the post administration urines as parent compound besides the metabolite desoxy-ecdysterone. Additionally desoxy-poststerone was tentatively assigned as minor metabolite, however further investigations are needed.

**Conclusion:** An administration of ecdysterone can be targeted using existing procedures of anti-doping laboratories. Ecdysterone and desoxy-ecdysterone appeared as suitable candidates for integration in ITP. Using dilute-and-inject a detection of the parent compound was possible for more than two days after the administration of a single dose of ~50 mg.

## Introduction

Ecdysterone (chemical structure in Figure 1) is widely marketed as a “natural anabolic agent”, advertised to increase strength and muscle mass during resistance training, to reduce fatigue and to ease recovery. Growth promoting and anabolic effects in various animal species including humans have been reported [1–18]. Ecdysterone appeared to promote an anabolic effect that was reported to be even stronger than that of the anabolic androgenic steroid (AAS) metandienone [19,20]. Contrary to the effect of AASs, it appears that the effect of ecdysterone is mediated by an activation of estrogen receptor beta (ERbeta) [1,21–23]. A few studies reported a performance enhancing effect in animals, but only recently a controlled administration trial in humans showed significant performance enhancement in resistance training [24]. Thus, the administration of ecdysterone may be considered as a practice that leads to an unfair advantage in sports competitions and may therefore be considered as doping. Indeed, ecdysterone was already suspected to be used by Olympic athletes since the 1980s and was also called a “Russian secret” [11,19,20,24–26].

**Figure 1:**
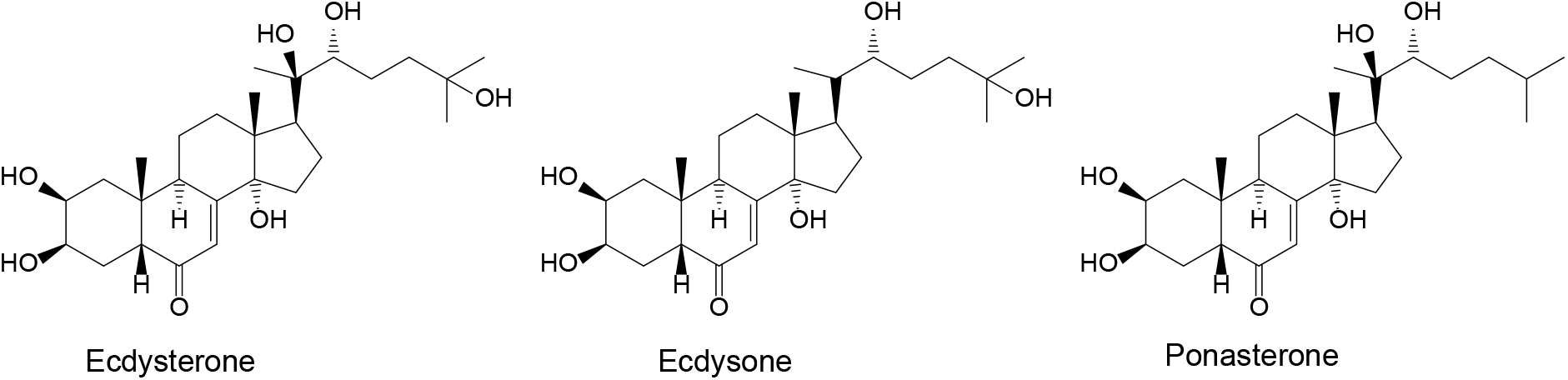
Chemical structure of ecdysterone, ecdysone (20-desoxy-ecdysone), and ponasterone (25-desoxy-ecdysone)

The administration of prohibited substances in sports is generally traced back from the analysis of biological specimen. The majority of substances is covered by the analysis of urine samples from athletes utilizing mass spectrometric detection hyphenated to chromatographic separation [27,28]. Sample preparation often includes cleavage of potential phase-II metabolites, concentration of the analytes using liquid-liquid (LLE) or solid phase extraction (SPE). For GC-MS analysis derivatization of the analytes to their TMS-derivatives is most frequently performed for less volatile compounds [28–30].

A few studies report the metabolism of ecdysterone, however most of them do not refer to humans [3,31–34]. As stated by Lafont et al. [31] considerable differences between species have been observed and thus, transferability is questionable. In humans, desoxy-ecdysterone was reported as urinary metabolite, however structure assignment differed between the two studies. While Brandt [34] reported 14-desoxy-ecdysterone, a 2-desoxy metabolite was described in Tsitsimpikou et al. [33].

Herein we investigate the analytical properties of ecdysterone and evaluate different alternatives to trace back its administration from the analysis of doping control samples using GC- or LC-MS based methods.

## Materials and methods

### Chemicals and reagents

Reference substances of the phytosteroids ecdysterone (2β,3β,14α,20β,22R,25-hexahydroxy-5β-cholest-7-en-6-one, parent compound, **PC**) and ecdysone (2β,3β,14α,22R,25-pentahydroxy-5β-cholest-7-en-6-one, **1**) were purchased from Steraloids (Newport, USA), while ponasterone (2β,3β,14α,20β,22R-pentahydroxy-5β-cholest-7-en-6-one, **2**) was obtained from Cayman Chemical Company (Ann Arbor, Michigan USA). Methyltestosterone (used as internal standard for urine analysis) was obtained from Sigma Aldrich (Milano, Italy).

The preparation of β-glucuronidase from Escherichia coli was from Roche Diagnostic (Mannheim, Germany). The derivatizing agent (TMIS reagent) was a mixture of N-methyl-N-trimethylsilyl-trifluoroacetamide (MSTFA)/ammonium iodide (NH_4_I)/dithierythritol (DTE) (1000:2:4 v/w/w) stored in screwed cap vials at 4 °C for a maximum of two weeks. MSTFA was supplied by Chemische Fabrik Karl Bucher GmbH (Waldstetten, Germany), NH_4_I and DTE were supplied by Sigma-Aldrich (Milano, Italy). Solvents (t-butyl methyl ether, ethyl acetate, methanol) and reagents (potassium carbonate, sodium phosphate, sodium hydrogen phosphate, sodium hydrogen carbonate) were of analytical or HPLC grade and provided by Sigma-Aldrich (Milano, Italy) or VWR (Darmstadt, Germany). Water was obtained from a MilliQ water purification system (Millipore S.p.A., Milano, Italy) or a LaboStar 2-DI/UV system (SG Wasseraufbereitung und Regeneration GmbH, Barsbüttel, Germany) equipped with LC-Pak Polisher and a 0.22-μm membrane point-of-use cartridge (Millipak).

### Synthesis of 14-desoxy-ecdysterone (3)

Desoxygenation of ecdysterone was performed as described by Kumpun et al. [3]. A solution of 200 mg of ecdysterone in 2.4 mL of acetic acid was treated with 250 mg of zinc powder while stirring for 24 h at 70°C. After filtration an aliquot was diluted with water and analyzed by LC-QTOF-MS. Three desoxy-ecdysterone isomers were detected with the two major assigned according to Kumpun et al. and Zhu et al. [3,35] to the 14α-H (most abundant, earlier eluting, **3**) and 14β-H product (second most abundant, later eluting, **3b**).

### Urine samples

Post administration urines were collected following the administration of 51.5 mg of ecdysterone (checked for identity and purity) to one healthy volunteer. All samples (as they accrued) were collected for 33 hours, followed by spot urines of the following three days. All samples were anonymized and handled in accordance with the ethical standards of the Helsinki Declaration. Informed consent form was signed by the volunteer. Samples were stored in aliquots at −18°C and gently thawed at +4°C for sample preparation.

### Instrumentation

Analyses were performed evaluating different instruments, i.e. GC-QTOF-MS, LC-QTOF-MS and LC-QQQ-MS. Mass Hunter B10 software from Agilent Technologies Inc. (Santa Clara, CA, USA) and Analyst Version 1.6.2 (AB Sciex, Monza, Italy) software were used for data acquisition and processing.

#### LC-QTOF-MS

LC-HRMS was performed on an Agilent 6550 Q-TOF mass spectrometer coupled to an Agilent 1290 II UHPLC equipped with an Agilent Eclipse Plus C18 column (2.1 mm x 100 mm, particle size 1.8 μm). A linear gradient (starting at 5% B and increasing to 25% at 7 min, then to 95% at 13 min, 3 min hold, followed by 2 min re-equilibration at 5% B) was used with water containing formic acid (1000:1, v:v, eluent A) and acetonitrile and formic acid (1000:1, v:v, eluent B) as mobile phase constituents at a flow rate of 0.4 mL/min. Ionization was performed by electrospray ionization (ESI) in positive mode using a Jet Stream ESI source and Ion Funnel (Agilent Technologies GmbH, Waldbronn, Germany). In ESI+ a capillary voltage of 3,500V, a nozzle voltage of 300 V, a drying gas flow of 15 L/min at 150°C, sheath gas flow of 12 L/min at 375°C and a nebulizer pressure of 25 psi were used. The high-pressure ion funnel was operated at radio frequency (RF) voltage 150 V, the low-pressure funnel at RF 60 V, and the octopole at RF 750 V.

The analyses were performed in full scan MS as well as targeted MS/MS mode or auto MS/MS mode at a mass range of m/z 50-1000. In MS/MS experiments nitrogen was used as collision gas and collision energies were ramped with precursor mass (slope: 6; offset: 4). Mass resolution (full width at half maximum, FWHM) within the analyzed m/z range was 12,000 to 25,000 (based on the QTOF mass calibration data of 118.086255 to 1521.971475). Purine ([M+H]^+^=121.0509 and the Agilent compound HP0921 ([M+H]^+^=922.0098) were simultaneously introduced into the ion source and used for internal mass calibration throughout the analysis.

#### LC-QQQ-MS

The LC-MS/MS instrument comprised of an Agilent 1200 with binary gradient system (Agilent Technologies S.p.A, Cernusco sul Naviglio, Milano, Italy) coupled to an API 4000 triple quadrupole mass spectrometer (ABSciex, Monza, Italy). Ionization was performed by ESI in positive mode, using a curtain gas pressure of 25 psi, a source temperature of 550 °C, an ion source gas 1 pressure of 35 psi, an ion source gas 2 pressure of 40 psi, a declustering voltage of 80 V, an entrance potential of 10 V and a needle voltage of 5500 V. The experiments were performed using multiple reaction monitoring (MRM) as acquisition mode, employing collision-induced dissociation (CID) using nitrogen as collision gas at 5.8 mPa, obtained from a dedicated nitrogen generator system Parker-Balston model 75-A74, gas purity 99.5% (CPS analitica Milano, Italy). The ion transitions selected for ecdysterone were: 481→371 (CE=28 eV), 481→445 (CE=25 eV), and 481→165 (CE=35 eV), whereas for ecdysone and ponasterone the selected ion transitions were 465→447 (CE=25 eV), 465→429 (CE=27 eV) and 465→285 (CE=27 eV).

The RP-HPLC separation was adopted from the method currently used in the World Anti-Doping Agency (WADA) accredited anti-doping laboratory of Rome [36] using a Supelco discovery C18 column (2.1 x 150mm, 5 μm) with a mobile phase of water containing formic acid (1000:1, v:v, eluent A) and acetonitrile and formic acid (1000:1, v:v, eluent B). The gradient program starts at 10% of eluent B and increases to 60% of eluent B in 3 min, 6 min hold, then to 90% of eluent B in 2 min, 1 min hold, followed by 4 min re-equilibration at 10% of eluent B. The flow rate was set at 250 μL/min and the column was maintained at ambient temperature. Aliquots of 10 μL were injected using an Agilent 1200 autosampler.

#### GC-QTOF-MS

GC-QTOF-MS was performed as regularly done in the Anti-Doping Laboratory in Rome. An Agilent 7200 GC-QTOF-MS (Waldbronn, Germany) coupled by electron ionization (EI, 70 eV at 250 °C) to an Agilent 7890B GC was equipped with an HP1 Ultra1 capillary column (length 17 m, 0.2 mm i.d., 0.11 μm film thickness). Helium was used as carrier gas (1 mL/min, constant flow) and the oven temperature was programmed at 188 °C (2.5 min hold), was increased at 3 °C/min to 211 °C (2 min hold), at 10 °C/min to 238 °C, at 40 °C/min to 320 °C (hold 3.2 min). An aliquot of 2 μL was injected in split mode (1:20). Improved separation used a modified temperature program, however, evaluation was intended to use the routinely performed separation of the laboratory to meet initial testing procedures.

Reference solutions or the final extracts of sample preparation were evaporated and derivatized using 100 μL of TMIS reagent.

### Evaluation of pre-analytical sample processing

#### Extraction procedures

To evaluate the possibilities of extraction and concentration of the analytes, ecdysterone parent compound was used as model substance. For solid phase extraction different cartridges (HLB Oasis HLB and WCX, both Waters, Milan, Italy, as well as Chromabond C18, C18 Hydra, and HR-X, all Macherey-Nagel, Düren Germany), were tested. After conditioning, cartridges were loaded with 2 mL of urine spiked with ecdysterone. After washing elution was performed with 1 mL of methanol (either with or without addition of formic acid or ammonium formate). The eluate was evaporated and reconstituted in 50 μL of purified water to yield the injection solution for LC-MS analysis.

Additionally, liquid-liquid extraction was evaluated at different pH values and using different extraction solvents. Aliquots of 2 mL of blank urine spiked with ecdysterone were adjusted to pH 5, 7 or 9 and extracted with 5 mL of solvent (TBME, ethyl acetate, chloroform, or a mixture of chloroform with isopropanol). Analyses were performed by LC-MS/MS.

Recoveries were determined by comparison with a reference solution spiked into urine after the extraction.

### Application to post administration samples

For LC-MS analysis urine samples were spiked with the internal standard methyltestosterone, diluted with water (1:4, v:v) and injected into the system after centrifugation at 800 *g*.

Alternatively, GC-QTOF analysis was performed after extraction and derivatization using a mixture of MSTFA:NH_4_I:DTE (1000:2:4, v/w/w, TMIS reagent) within 2 h at 75 °C.

For evaluation of a potential cleavage of phase-II metabolites enzymatic hydrolysis with β-glucuronidase or a mixture of β-glucuronidase and arylsulfatase was performed before the analysis.

To 2 mL of each urine sample adjusted to pH 7 by addition of 1.5 mL of phosphate buffer, 50 μL of internal standard solution (methyltestosterone, 200 ng/mL) and 50 μL of β-glucuronidase from *Escherichia coli* (Roche Diagnostics, Mannheim, Germany) were added. Deglucuronidation was performed within 1 hour at 50 °C.

Additionally cleavage of potential glucuronides and sulfates was performed while incubating 2 mL of urine, adjusted to pH 5.2, with 50 μL of β-glucuronidase arylsulfatase mix after addition of the internal standard (methyltestosterone). Incubation was carried out for 3 h at 50°C.

## Results and discussion

### Mass spectrometry

The mass spectrometric properties of ecdysterone and the two other ecdysteroids ecdysone and ponasterone (both desoxy analogues of ecdysterone accessible as reference materials) as well as the synthesized 14-desoxy-ecdysterone were analyzed by GC-MS and LC-MS. Derivatization, that was required for GC-MS analyses, was performed using TMIS reagent that is most frequently utilized in anti-doping laboratories. Ecdysterone mainly yielded a per-TMS derivative (ecdysterone 5-en-6-ol heptakis-TMS derivative) eluting at 18.69 min. The most abundant fragments were detected at m/z 633.3593 (C_33_H_61_O_4_Si_4_^+^, exact mass 633.3641, mass error Δm/z=6.6 ppm), m/z 543.3094 (C_30_H_51_O_3_Si_3_^+^, exact mass 543.2621, mass error Δm/z=8.3 ppm), and m/z 171.1189 (C_9_H_19_OSi^+^, exact mass 171.1200, mass error Δm/z=4.7 ppm). Similar findings were reported by Tsitsimpikou et al. using GC-MS with single quadrupole [33]. The EI spectrum including proposed generation of fragments is displayed in Figure 2. Additionally, Figure 2 displays the product ion spectrum obtained from LC-ESI-QTOF-MS analysis. In MS 1 analysis ecdysterone was detected as [M+H]^+^=481.3170 (C_27_H_45_O_7_^+^, exact mass 481.3160, mass error Δm/z=2.1 ppm) that may also face 1-3 water losses as in-source fragmentation. MS/MS of the molecular ion as precursor resulted in major fragments at m/z 371.2209, m/z 445.2933, and m/z 165.1275 that were also used for MRM in LC-QQQ-MS analysis. While m/z 445.2933 (C_27_H_41_O_5_^+^, exact mass 445.2949, mass error Δm/z=3.6 ppm) results from the loss of two water molecules from [M+H]^+^, m/z 371.2209 is most likely generated from an α-cleavage of the C23-C24 bond in the side chain and an additional water loss resulting in C_23_H_31_O_4_^+^ (exact mass 371.2217, mass error Δm/z=2.2 ppm). M/z 165.1275 (C_11_H_17_O^+^, exact mass 165.1274, mass error Δm/z=0.6 ppm) may be explained by a fragment including C15-C17 from the D-ring including the attached side chain after losses of two water molecules, while the side chain (C20-C27) after the loss of two water molecules results in m/z 125.0961 (C_8_H_13_O^+^, exact mass 125.0961, mass error Δm/z=0 ppm). Both explanations are in-line with the spectrum of makisterone (24-methyl-ecdysterone), which displays an analogous fragments at m/z 179 and m/z 139 [34].

**Figure 2:**
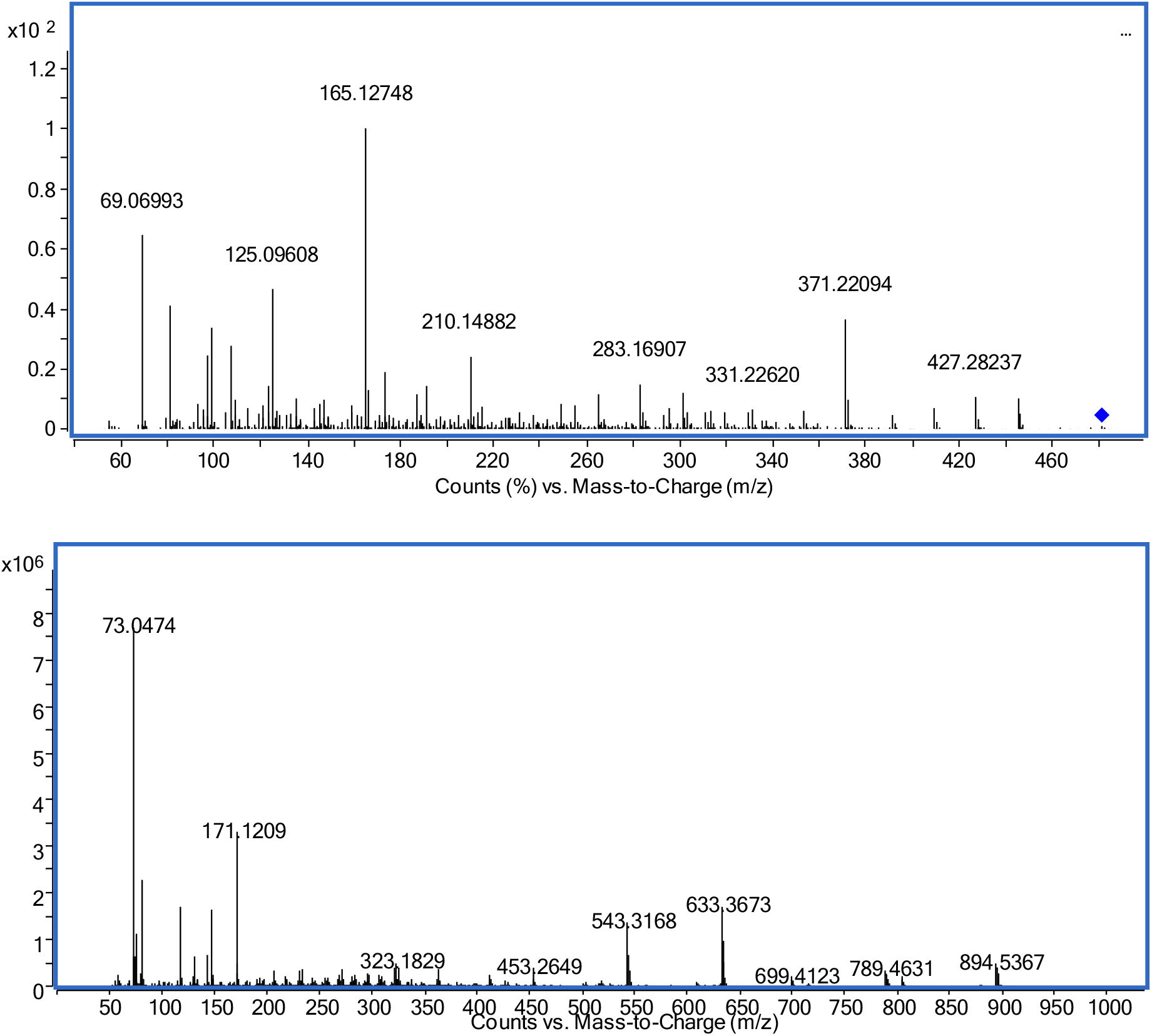
Mass spectra of ecdysterone; upper: product ion spectrum LC-QTOF-MS (precursor [M+H]^+^_theor_=481.3160, RT=5.80 min); lower: GC-EI-QTOF-MS as per-TMS (M_theor_^+•^=984.5848, RT=18.69 min)

LC-QTOF-MS analysis of ecdysone resulted in [M+H]^+^=465.3209 (C_27_H_45_O_6_^+^, exact mass 465.3211, mass error Δm/z=0.4 ppm) even dominated by [M+H-H_2_O]^+^=447.3118 (C_27_H_43_O_5_^+^, exact mass 447.3105, mass error Δm/z=2.9 ppm). MS/MS of the molecular ion as well as [M+H-H_2_O]^+^ as precursor (Figure 3) results in major fragments at m/z 429.2997 ([M+H-H_2_O]^+^, C_27_H_41_O_4_^+^, exact mass 429.2999, mass error Δm/z=0.6 ppm), and m/z 109.1013 (C20-C27 after loss of two water molecules as analogously described for ecdysterone, C_8_H_13_^+^, exact mass 109.1012, mass error Δm/z=1.5 ppm). Similarly, ponasterone yielded [M+H]^+^=465.3221 (C_27_H_45_O_6_^+^, exact mass 465.3211, mass error Δm/z=2.0 ppm) with fragmentation dominated by water losses ([M+H-H_2_O]^+^=447.3088, C_27_H_43_O_5_^+^, exact mass 447.3105, mass error Δm/z=7.6 ppm, [M+H-2*H_2_O]^+^=429.2984, C_27_H_41_O_4_^+^, exact mass 429.2999, mass error Δm/z=5.9 ppm, [M+H-3*H_2_O]^+^=411.2890, C_27_H_39_O_3_^+^, exact mass 411.2894, mass error Δm/z=1.3 ppm, [M+H-4*H_2_O]^+^=393.2807, C_27_H_37_O_2_^+^, exact mass 393.2788, mass error Δm/z=1.4 ppm). As described for ecdysone m/z 109.1009 may be explained as fragment C20-C27 after loss of two water molecules, C_8_H_13_^+^, exact mass 109.1012, mass error Δm/z=2.7 ppm). The MS/MS spectrum is displayed as Figure 4.

**Figure 3:**
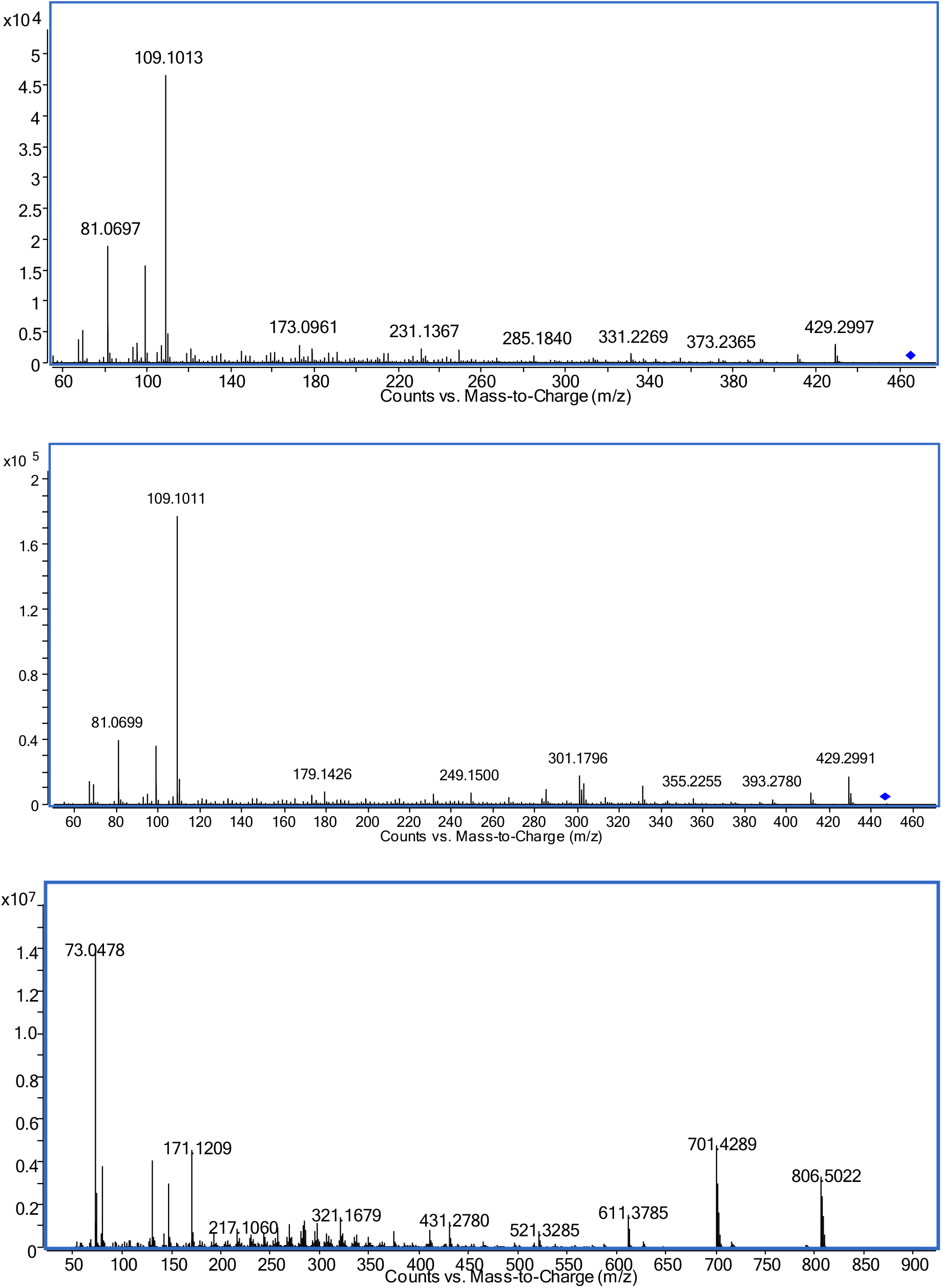
Mass spectra of ecdysone; upper: product ion spectrum LC-ESI-QTOF-MS (precursor [M+H]^+^_theor_=465.3211, RT=7.12 min); middle: product ion spectrum LC-ESI-QTOF-MS (precursor [M+H-H_2_O]^+^_theor_ =447.3105, RT=7.12 min); lower: GC-EI-QTOF-MS as per-TMS (M_theor_^+•^=896.5504, RT=18.12 min)

**Figure 4:**
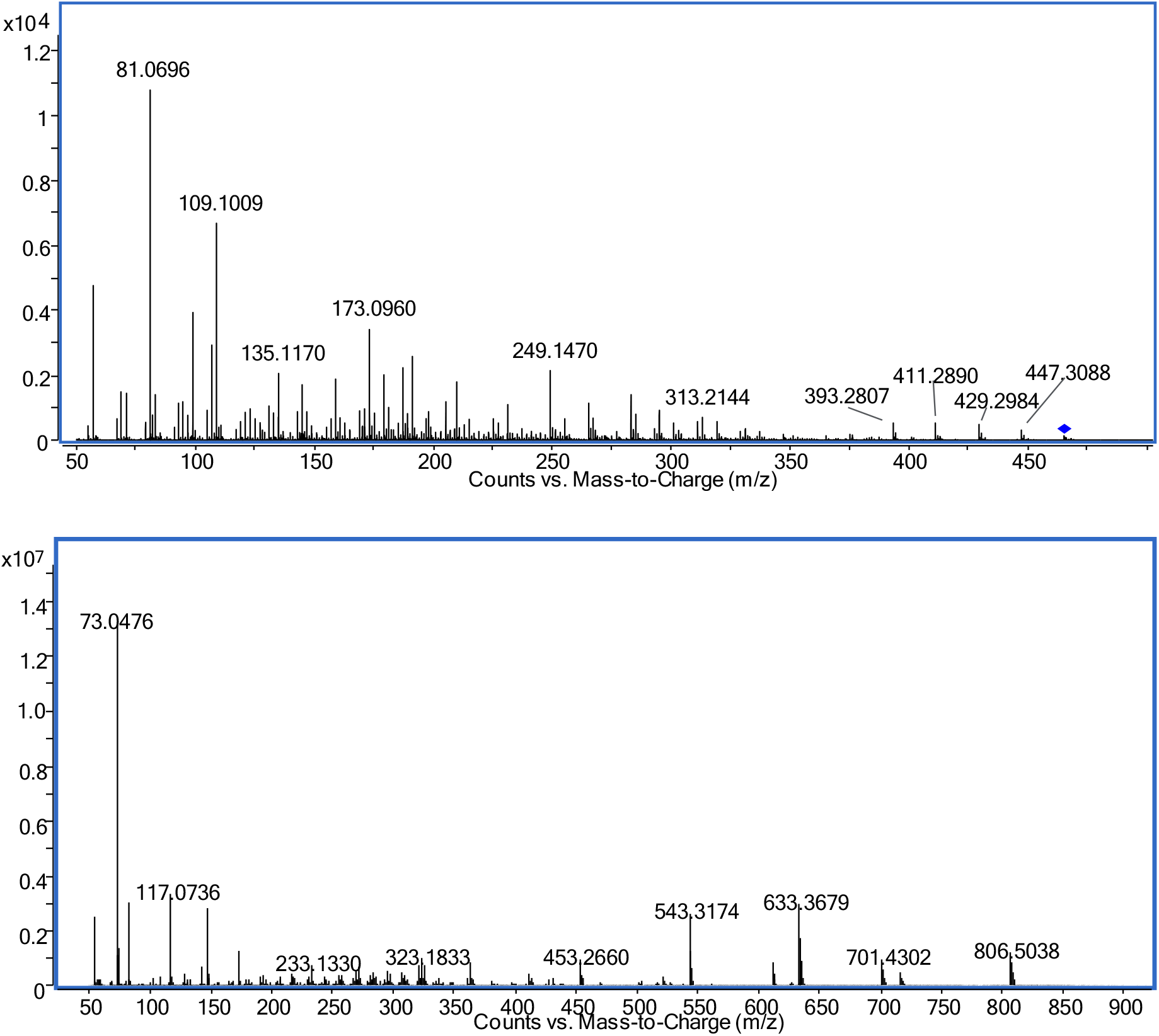
Product ion spectrum (LC-ESI-QTOF-MS) of ponasterone (precursor [M+H]^+^_theor_=465.3211, RT=8.85 min); lower: GC-EI-QTOF-MS as per-TMS (M_theor_^+•^=896.5504, RT=18.11 min)

Equally, the synthesized 14-desoxy-ecdysterone **(3)** was detected at [M+H]^+^=465.3209 (C_27_H_45_O_6_^+^, exact mass 465.3211, mass error Δm/z=0.4 ppm). The product ion spectrum (LC-QTOF-MS, Figure 5) of [M+H]^+^ yielded m/z 303.1956, which may be explained as fragment after full cleavage of the side chain, i.e. C1-C19 (C_19_H_27_O_3_^+^, exact mass 303.1955, mass error Δm/z=0.3 ppm). One or two losses of water out of this result in m/z 285.1850 (C_19_H_25_O_2_^+^, exact mass 285.1849, mass error Δm/z=0.4 ppm) and m/z 267.1743 (C_19_H_23_O^+^, exact mass 267.1743, mass error Δm/z=0 ppm). As described for ecdysterone m/z 125.0962 is considered as fragment of the side chain after the loss of two water molecules (C_8_H_13_O^+^, exact mass 125.0961, mass error Δm/z=0.8 ppm). Furthermore, m/z 191.1068 may be assigned to C1-C11+C19 after loss of water (C_12_H_15_O_2_^+^, exact mass 191.1067, mass error Δm/z=0.5 ppm). Similar fragments were reported by Brandt [34], however, in unit mass resolution.

**Figure 5:**
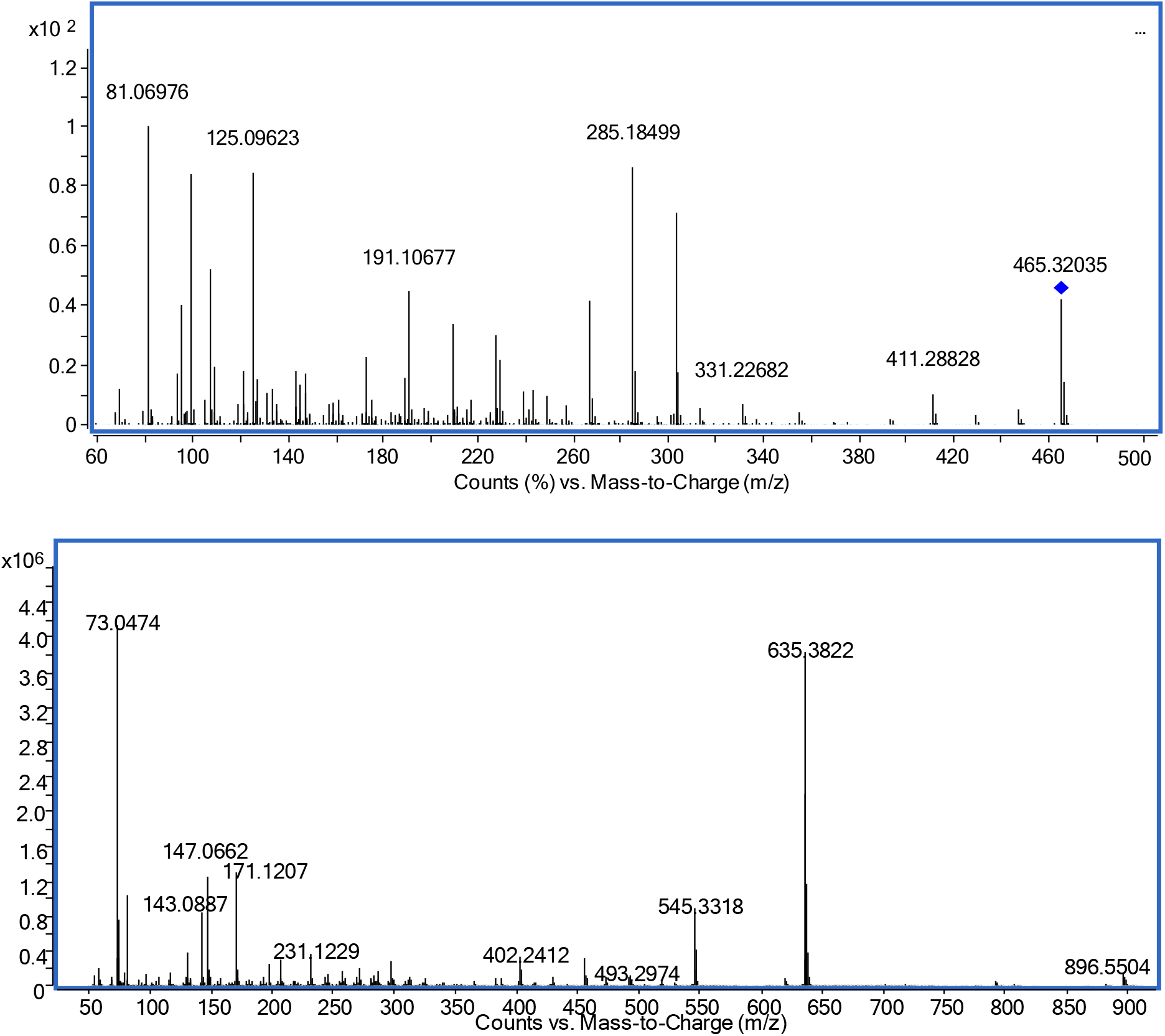
Product ion spectrum (LC-ESI-QTOF-MS) of 14-desoxy-ecdysterone (precursor [M+H]^+^_theor_=465.3211, RT=6.86 min); lower: GC-EI-QTOF-MS as per-TMS (M_theor_^+•^=896.5504, RT=18.65 min)

Following derivatization 14-desoxy-ecdysterone **(3)** yielded a per-TMS derivative as well (14-desoxy-ecdysterone 5-en-6-ol hexakis-TMS derivative) eluting slightly earlier than ecdysterone-per-TMS (RT_**M3**_ = 18.65 min). The most abundant fragments were detected at m/z 635.3822 (C_33_H_63_O_4_Si_4_^+^, exact mass 635.3798, mass error Δm/z=3.8 ppm), m/z 545.3318 (C_30_H_53_O_3_Si_3_^+^, exact mass 545.3297, mass error Δm/z=3.8 ppm), and m/z 171.1207 (C_9_H_19_OSi^+^, exact mass 171.1200, mass error Δm/z=4.1 ppm).

The retention times of the above mentioned reference steroids in LC-QTOF-MS are listed in Table 1.

**Table 1:** Retention times and [M+H]^+^ of ecdysteroids analyzed by LC-ESI-QTOF-MS

| No | Name | RT [min] | $[M+H]^+_{\text{theor}}$ |
| --- | --- | --- | --- |
| PC | ecdysterone | 5.80 | 481.3160 |
| 1 | ecdysone | 7.12 | 465.3211 |
| 2 | ponasterone | 8.85 | 465.3211 |
| 3 | 14-desoxy-ecdysterone | 6.86 | 465.3211 |
| M3 | urinary desoxy-ecdysterone | 6.92 | 465.3211 |
| M4 | urinary 14-desoxy-poststerone isomer 1 | 8.84 | 347.2217 |
| M5 | urinary 14-desoxy-poststerone isomer 2 | 9.36 | 347.2217 |

### Extraction procedure

Ecdysterone is known as a relatively polar steroid (logP = −0.53, according to ACDLabs prediction and reported at www.scifinder.cas.org). Ecdysone and ponasterone show slightly less polarity (logP_ecdysone_ = 0.875, logP_ponasterone_ = 1.434), thus still even more polar compared to cortisol (logP_cortisol_ = 1.762, for comparison logP_testosterone_ = 3.179). Extraction was therefore evaluated using TBME that is utilized in lots of extraction procedures in anti-doping laboratories and compared with less common solvents such as ethyl acetate, chloroform, or a mixture of chloroform with isopropanol. Almost independent of the pH of the aqueous phase extraction yields in LLE were generally low (<23%) with TBME giving the worst recovery (<5%) of all extraction solvents tested in this investigation. SPE using Oasis HLB resulted in the highest recovery (96%) for extraction after optimization of the procedure. According to literature Lichrolut RP-18 also gave recoveries of 72-96% [34]. Thus, it is suggested to perform either dilute-and-inject or SPE for sample preparation rather than LLE.

### Detection of ecdysterone metabolites in urine

Considering the above mentioned pre-analytical properties it was decided to investigate the urinary elimination of ecdysterone using dilute-and-inject. The majority of experiments was performed using LC-QTOF-MS in MS1 mode, with MS/MS experiments utilized for confirmation. In the post administration urines the parent compound ecdysterone was found as most abundant analyte. No differences in abundance were found after enzymatic hydrolysis either with β-glucuronidase or β-glucuronidase/arylsulfatase. Additionally, neither ecdysterone glucuronide nor ecdysterone sulfate were detected in the non-hydrolyzed samples. This suggests an excretion in unconjugated form. Furthermore, the presence of a desoxy metabolite was confirmed in the post administration urines. It was detected in MS1 ([M+H]^+^=465.3202, C_27_H_45_O_6_^+^ exact mass 465.3211, Δm/z=1.9 ppm). 20-Desoxy- and 25-desoxy-ecdysterone were excluded by comparison with the authentical references of ecdysone and ponasterone. Utilizing MS/MS (spectrum in Figure 6 upper), the comparison with the product ion spectrum of 14-desoxy-ecdysterone as reported in Figure 5 revealed high similarities, thus, considering high probability of conformance. Additionally, the GC-QTOF-MS analysis showed a signal matching 14-desoxy-ecdysterone spectrum and retention time. Fragments look very similar to those reported for 2-desoxy-ecdysterone by Tsitsimpikou et al. [33], however showing different relative abundances. As already commented by Lafont and Dinan [10] mass spectrometry is not capable of providing data suitable for unambiguous assignment of the exact isomer. Even worse, also product ion spectra of ecdysteroids after ESI are dominated by losses of water that are not suitable for a discrimination of the position of desoxygenation at the sterane moiety. Similarly the EI-spectra reported by Tsitsimpikou et al. [33] are dominated by cleavage of the sterane moiety from the side chain and subsequent losses of TMSOH, that, again, do not provide suitable discrimination of the positional isomers. Thus, in the future synthesis of additional, authentic reference material is needed for proper isomer identification of isolated metabolites, by comparison of the retention times. As already discovered for ecdysterone, no relevant phase-II metabolite was detectable by accurate mass in the non-hydrolyzed samples and no increase in abundance was detected after enzymatic hydrolysis. This is in line with the findings in mice reported by Lafont et al. [31].

**Figure 6:**
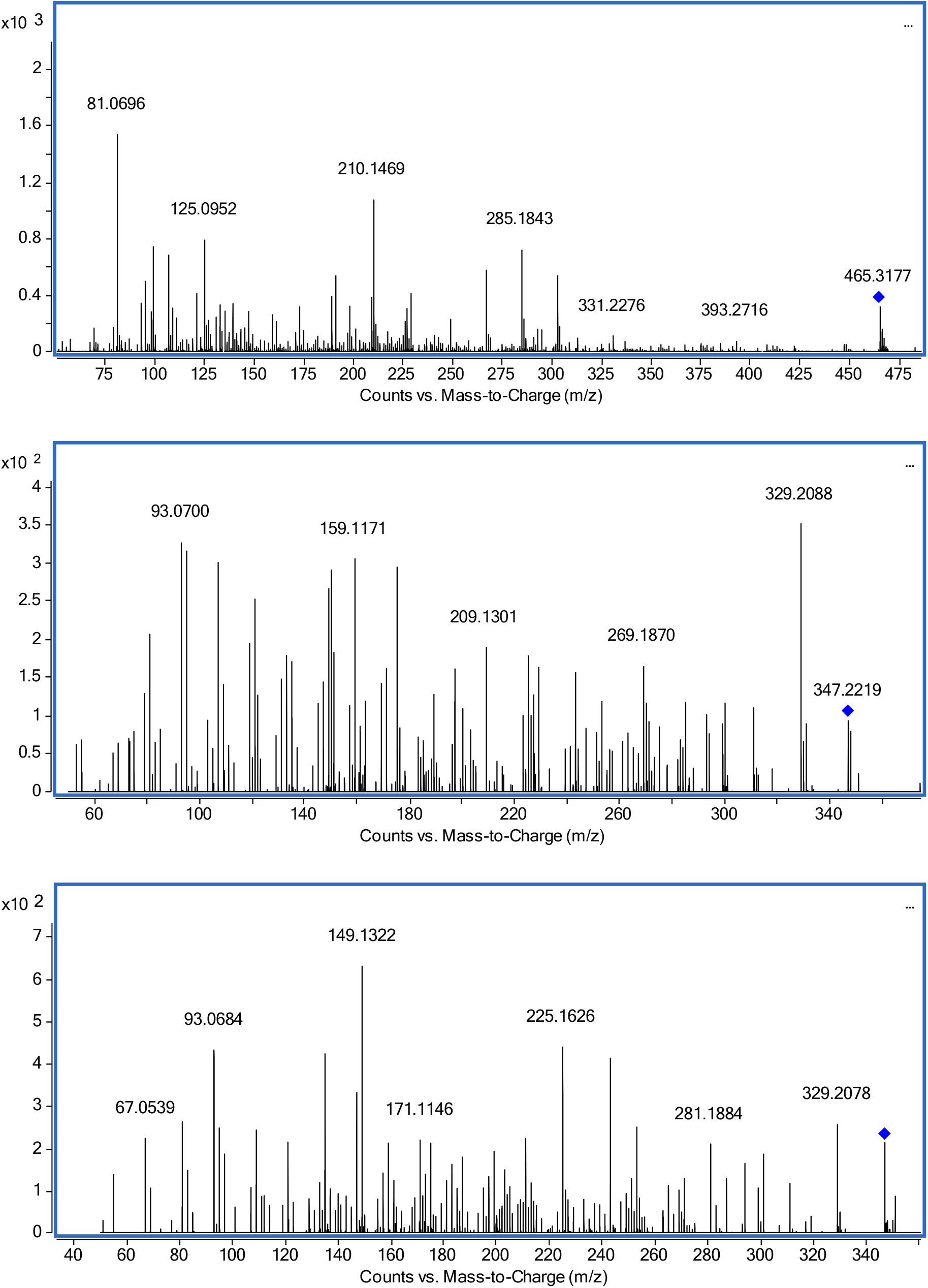
Product ion spectrum (LC-ESI-QTOF-MS) of urinary metabolites; upper: desoxy-ecdysterone (**M3**, precursor [M+H]^+^_theor_=465.3211, RT_**M3**_=6.92 min), middle and lower: two additional metabolites (**M4** and **M5**) tentatively assigned to desoxy-poststerone (precursor [M+H]^+^_theor_=347.2235, RT_**M4**_=8.84 min, RT_**M5**_=9.36 min)

Additionally Kumpun et al. [3] report the formation of poststerone (2β,3β,14α-trihydroxy-5β-pregn-7-ene-6,20-dione, C_21_H_30_O_5_, [M+H]_theor_^+^=363.2166) and 14-desoxy-poststerone (2β,3β-dihydroxy-5β-pregn-7-ene-6,20-dione, C_21_H_30_O_4_, [M+H]_theor_^+^=347.2217) in mice based on comparison with reference substances. Data mining based on this resulted in the detection of **M4** and **M5**, both compounds that may be assigned to desoxy-poststerone isomers (C_21_H_30_O_4_, [M+H]_theor_^+^=347.2217). Their product ion spectra are displayed in Figure 6. No signal that may correspond to poststerone (C_21_H_30_O_5_, [M+H]_theor_^+^=363.2166) was detected in our study. The structures of the above-mentioned metabolites reported in literature are summarized in Figure 7.

**Figure 7:**
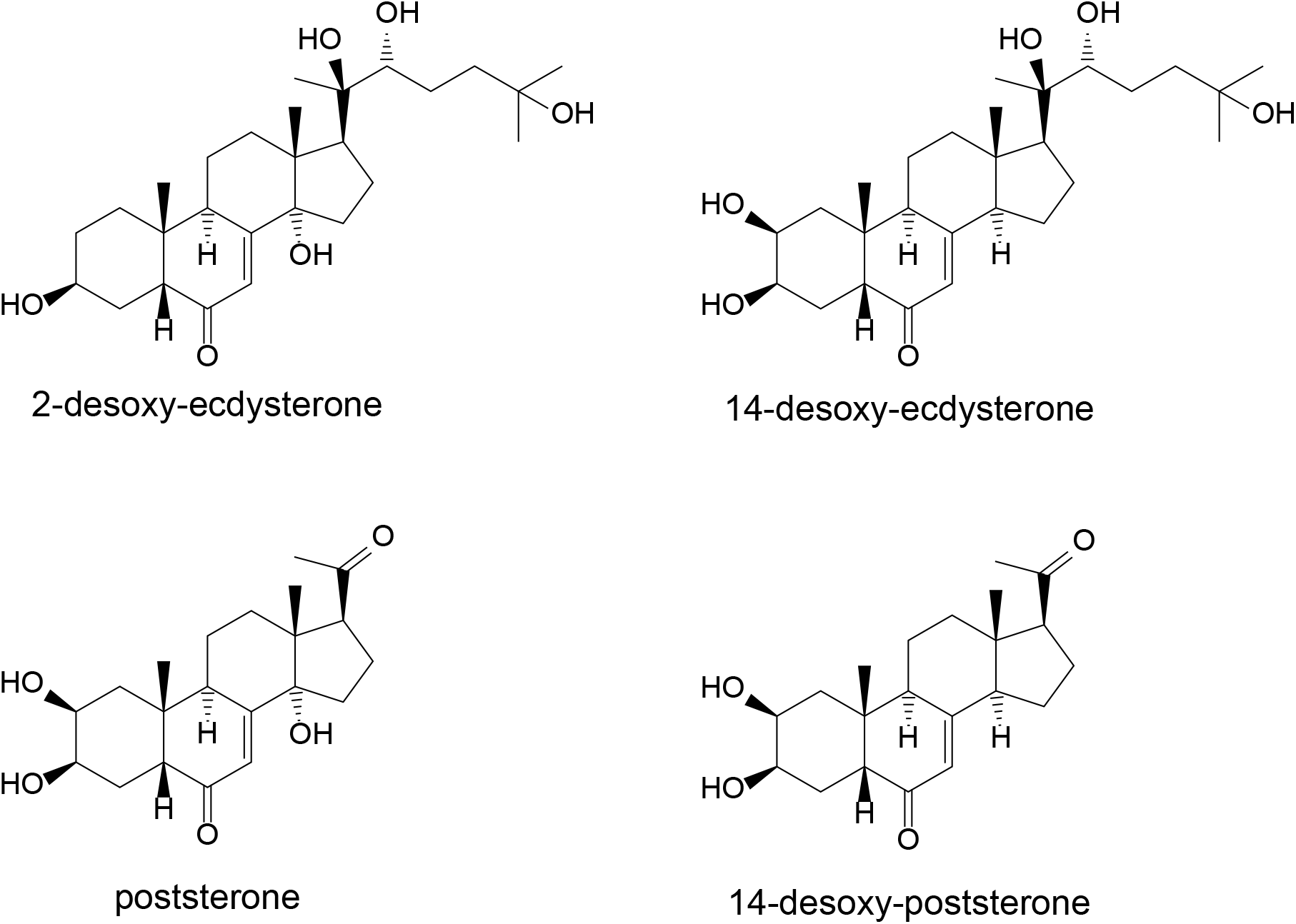
Chemical structures of desoxy-ecdysterone, poststerone and desoxy-poststerone proposed as metabolites of ecdysterone in literature [3,33,34]

Further metabolites described by Kumpun et al. [3] with [M+H]^+^=349 (two isomers), [M+H]^+^=351, and [M+H]^+^=365 were reported as reduced derivatives of poststerone or 14-desoxy-poststerone therein. The structures of the latter were tentatively assigned based on MS data. Only the dihydro-ecdysone isomer 5β-cholest-8(14)-ene-2β,3β,6α,22R,25-pentol) was reported to be substantiated by NMR and UV as well. None of these compounds was detected in our study. As the study of Kumpun et al. was performed in mice with the analysis accomplished based on urine and feces with no clear indication which metabolite was excreted in which specimen, the transfer to our study may be not fully applicable. Pathways of metabolite generation should be evaluated in the future to better understand potentially influencing factors. If desoxygenation is indeed caused by gut bacteria as postulated by Kumpun et al. [3] the bacterial composition may highly influence metabolite excretion. An evaluation of post administration urines of different volunteers is thus highly desired.

### Excretion profiles

Following a single oral dose of 51.5 mg of ecdysterone the parent compound was detectable by LC-QTOF-MS for more than two days using dilute-and-inject. The maximum concentration was observed in the 2.0-3.5 h urine. The desoxy metabolite M3 shows a smaller detection window compared to the parent compound. The maximum concentration was detected in the 7.25-9.0 h urine, and latest detection was achieved in the 25.2-29 h urine. The excretion profiles of both metabolites are displayed in Figure 8. A biphasic excretion of the two desoxy-poststerone isomers has been also observed, however needs further investigation.

**Figure 8:**
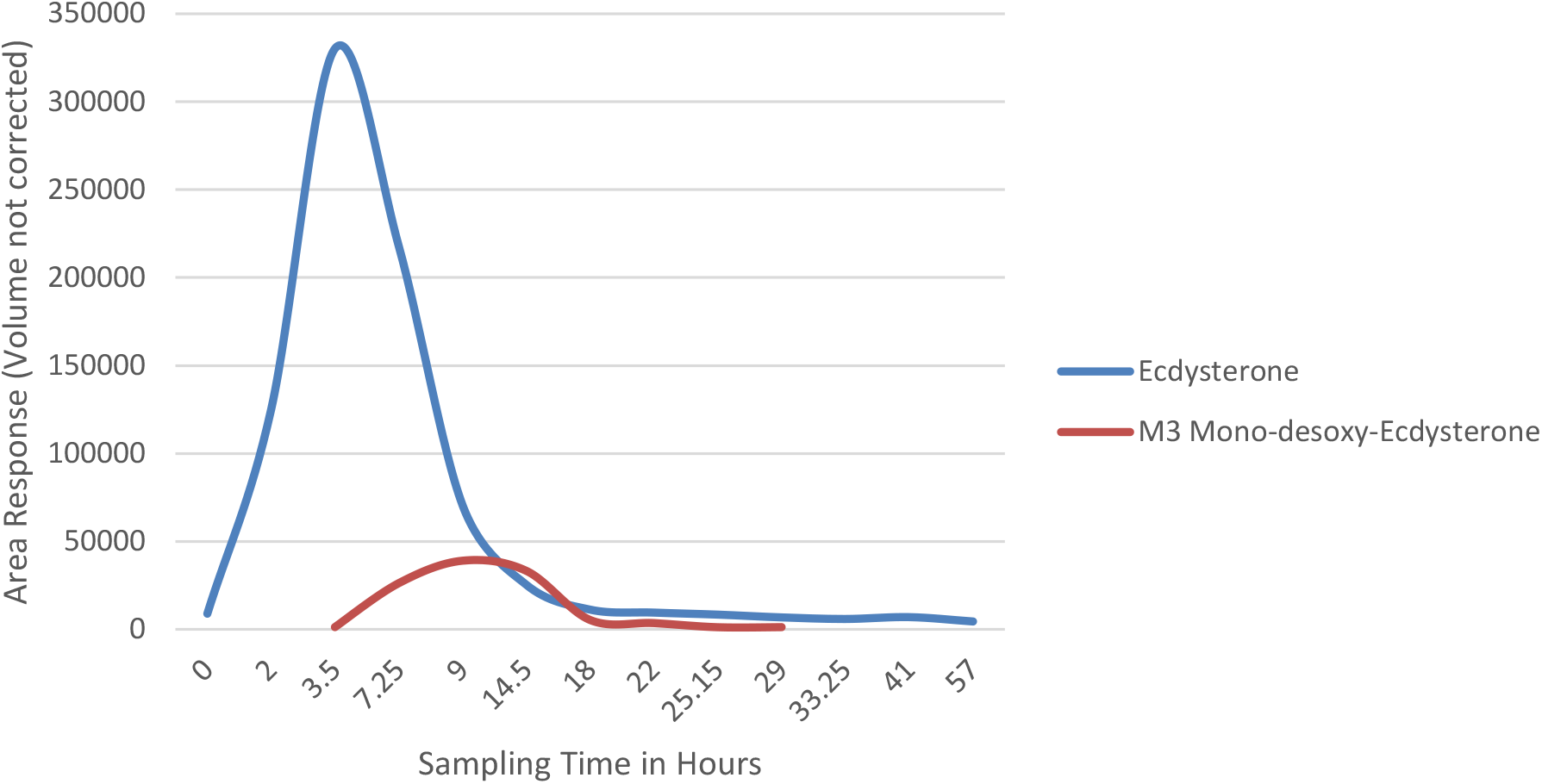
Excretion profiles (LC-ESI-QTOF-MS) of urinary ecdysterone and desoxyecdysterone (**M3**)

## Conclusion

An administration of ecdysterone results in an excretion of the parent compound in the urine. Targeting the parent compound is ideally performed by LC-HRMS or LC-MS/MS by dilute-and-inject. Alternatively, GC-MS analysis is possible after TMS derivatization. If extraction is required or desired, SPE was found by far superior to LLE. Enzymatic hydrolysis did not provide advantages over the analysis of the unconjugated fraction only. Ecdysterone and its desoxy metabolite may be easily integrated in current initial testing procedures (ITP) for monitoring the prevalence in elite sports.

Due to the potential generation of metabolites by gut bacteria that may cause significant variations in the metabolic profile, an integration of further isomers and analogues may also be appropriate. Furthermore ecdysterone is often administered from plant extracts such as spinach or suma root. Thus, further phytosteroids (e.g. ecdysone, ponasterone, and others) may be ingested aside. These steroids as well as their metabolites may be found in excretion urines as well.

## Acknowledgements

The World Anti-Doping Agency is acknowledged for their financial support (research grant 18C18MP). Mrs. Maxi Wenzel, Freie Universitaet Berlin, is acknowledged for technical assistance.

## Compliance with Ethical Standards

Financial support of the study was granted by the World Anti-Doping Agency. The authors declare no other conflict of interest.

